# ProNetView-ccRCC: A web-based portal to interactively explore clear cell renal cell carcinoma proteogenomics networks

**DOI:** 10.1101/2020.03.12.981357

**Authors:** Selim Kalayci, Francesca Petralia, Pei Wang, Zeynep H. Gümüş

## Abstract

To better understand the molecular basis of cancer, the NCI’s Clinical Proteomics Tumor Analysis Consortium (CPTAC) has been performing comprehensive large-scale proteogenomic characterizations of multiple cancer types. Gene/protein regulatory networks has subsequently been derived based on these proteogenomic profiles, which serve as useful tools to gain system-level understanding of the molecular regulatory factories underlying the diseases. On the other hand, it remains a challenge to effectively visualize and navigate the resulting network models, which capture higher order structures in the proteogenomic profiles. There is a pressing need to have a new open community resource tool for intuitive visual exploration, interpretation and communication of these gene/protein regulatory networks by the cancer research community. In this work, we introduce ProNetView-ccRCC (http://ccrcc.cptac-network-view.org/), an interactive web-based network exploration portal for investigating phosphopeptide co-expression network inferred based on the CPTAC clear cell renal cell carcinoma (ccRCC) phosphoproteomics data. ProNetView-ccRCC enables quick, user-intuitive visual interactions with the ccRCC tumor phosphoprotein co-expression network comprised of 3,614 genes, as well as 30 functional pathway-enriched network modules. Users can interact with the network portal and can conveniently query for association between abundance of each phosphopeptide in the network and clinical variables such as tumor grade.

---

Recent advances in molecular profiling technologies^[1,2]^ enable the large-scale integrative proteomic and genomic (proteogenomic) studies of cancers. For example, the National Cancer Institute’s Clinical Proteomics Tumor Analysis Consortium (CPTAC) has recently performed comprehensive proteogenomic characterizations of tumor samples from breast,^[3,4]^ colon,^[5]^ ovarian,^[6]^ and kidney^[7]^ cancer patients. Integrative analyses based on rich proteogenomic profiles from these studies have greatly advanced our understanding of the molecular mechanisms underlying these cancers. Specifically, the co-expression networks^[8–10]^ derived from proteomics and phosphoproteomics data revealed the regulatory relationships between proteins and post-transcriptional modifications (PTMs), bringing new insights on complicated pathway interactions and dysfunctions that drive tumor initiation and progression^[3] [7]^.

While these protein/PTM co-expression networks contain rich information for identifying active pathways and oncogenic drivers, they are challenging to visually communicate due to their large dimensions and complicated topologies. Furthermore, these networks are best visually explored together with associated information between gene/protein activities and biological pathway annotations as well as clinical outcomes. To better understand and interpret the multiple layers of data provided in these networks, effective visual exploration tools that address these challenges are needed.

Current network data visual exploration tools^[11,12]^ are mainly used for visually communicating networks as static 2D images in print publications. While these provide advanced capabilities for visual customizations, they are not well suited for user-initiated interactive network explorations. Furthermore, as incorporating and exploring the meta-data associated with the CPTAC co-expression networks will make them readily available to the cancer research community, a centralized platform that is customized for CPTAC co-expression networks and associated meta-data is necessary. There is thus a pressing need to have a new open community resource tool for intuitive visual exploration, interpretation and communication of these gene/protein regulatory networks by the cancer research community.

To address these needs, we have developed a web-based interactive portal, ProNetView-ccRCC, which provides an intuitive and customized exploration environment for phosphopeptide co-expression network and other analysis results from CPTAC ccRCC study^[7]^. Its 3D network exploration interface enables quick and easy interactions with the ccRCC phosphopeptide co-expression network as well as 30 functional network modules and their associated meta-data. Furthermore, users can easily query for their proteins of interest, neighbors they are connected to, mapping peptides or pathways, at user-defined filters on clinical variables (stage, grade, age, gender, FDR cut-off). The portal is an open-access freely available resource at http://ccrcc.cptac-network-view.org/.

## Methods

ProNetView-ccRCC enables visual explorations of two network types: i) the phosphopeptide co-expression network of ccRCC tumors and ii) 30 different network modules derived based on the topology of the overall network. The network characterizes the co-expression patterns among 20,976 phosphopeptides in ccRCC and was derived using a random-forest based network construction algorithm^[10]^ based on phospho-peptide level data of 103 ccRCC samples. The network displays association among genes based on phosphopeptide level data with two genes being connected if at least two phosphopeptides mapping to the two genes are associated to each other. Based on the overall network topology, we identified sets of genes (i.e., network modules) tightly connected in the network. Specifically, network modules were derived by using Glay^[13]^ clustering algorithm. 30 modules containing at least 20 genes were identified and each was presented as a different module network. For each module network, pathway enrichment analysis was performed via Fisher Exact test and those overrepresented pathways were incorporated within the portal as associated meta-data^[7]^.

For intuitive exploration of these dense and complex networks, ProNetView-ccRCC enables interactive explorations in 3D. We implemented this capability by building on the Three.js framework, which provides an interface for WebGL. We calculated the network layouts using 3D layout algorithms within iCAVE ^[14,15]^, which is a desktop-based tool we have developed earlier for the stereoscopic and immersive exploration of networks. To handle various user interaction events in real time, we have utilized multiple client-side Javascript libraries (e.g. D3.js, dataTables.js, JQuery, etc.). For the styling of web interface elements, we have primarily utilized Bootstrap v3.3.7, integrated with some styling of our own. Since ProNetView-ccRCC utilizes only standard libraries and does not require the use of any additional plug-ins, the portal runs on all modern web browsers.

### Use Protocols

ProNetView-ccRCC landing page (http://ccrcc.cptac-network-view.org/) provides the global ccRCC tumor phosphoproteomics co-expression network (Figure 1A) and links it to functional network modules (Figure 1B). Users can interactively explore the large and dense topology of the whole ccRCC phosphopeptide co-expression network in 3D by rotating and zooming in/out the view. Six modules within this network (Figure 1B) were significantly associated with biological pathways^[7]^, as highlighted within the network using different coloring of nodes. Users can easily query for genes of their interest in the network. For example, one of the genes highlighted in the CPTAC ccRCC paper^[7]^ was ERG, a gene associated to VEGF response. By searching for the oncogene ERG (Figure 1C), the corresponding node is highlighted in red in the network exploration panel (Figure 1A) and it can easily be identified as member of the Angiogenesis enriched module network (Figure 1B). The query further returns the list of genes whose phosphopeptides were directly connected to ERG (i.e. co-expressed phosphopeptides) and provides the association of its corresponding phosphoprotein with tumor grade (p-value: 0.003, FDR: 0.118) and other phenotypes of interest (Figure 1C). Overall, these help understand the phosphopeptide network in the context of markers of user interest to help guide hypothesis generation for future studies.

**Figure 1.**
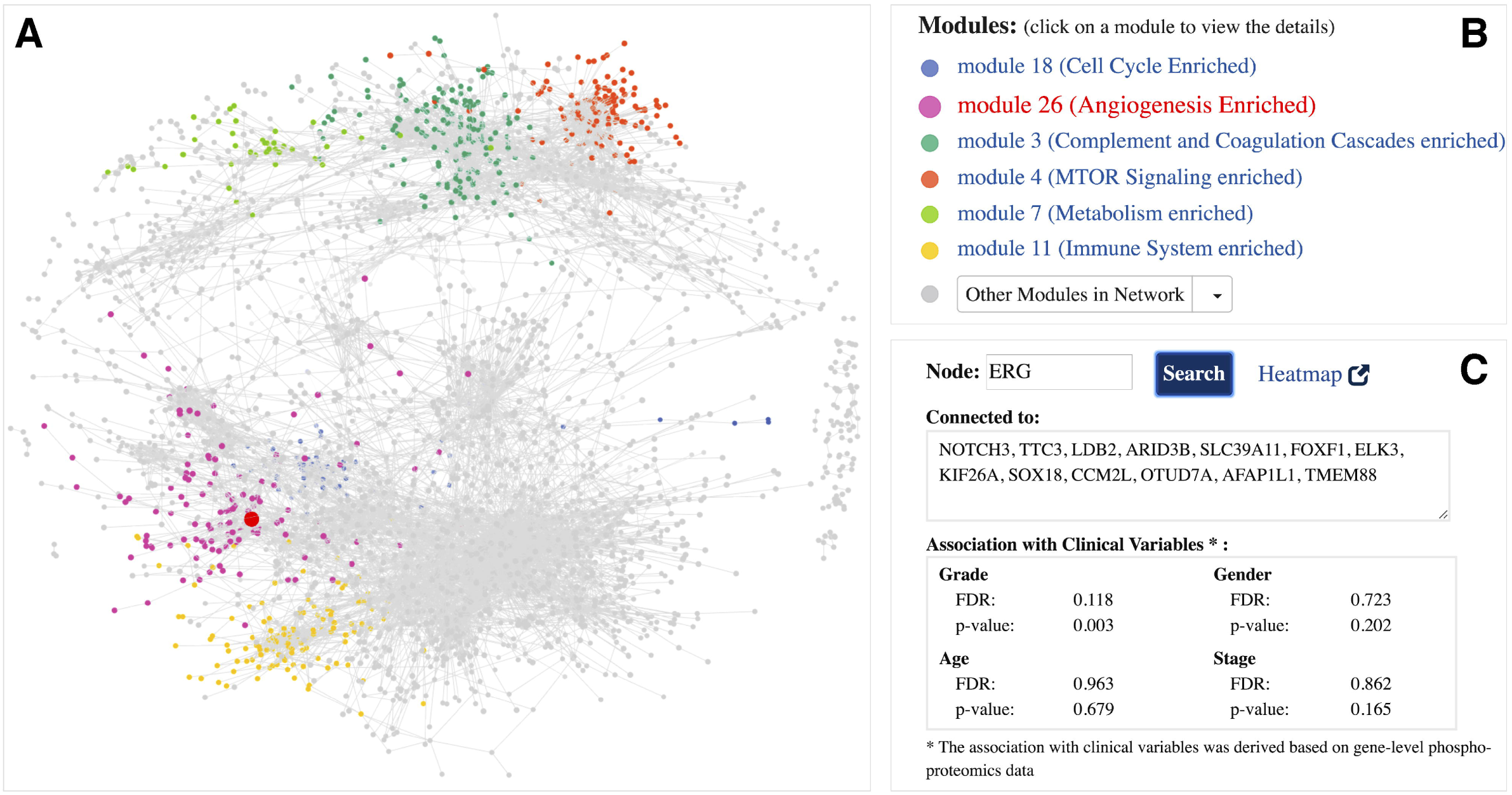
Global ccRCC tumor phosphoproteomics co-expression network **A** interactive 3D network exploration panel. Subsets of genes are color-coded based on their association with modules that are part of significant biological pathways. Large, red-colored node highlights the ERG gene; **B** list of module networks. 6 modules that are part of significant biological pathways are color-coded, the remaining 24 module networks are listed under the Other Modules in Network dropdown menu; **C** meta-data information associated with ERG gene. Gene-specific information includes the list of connections (co-expressed genes) and its association with clinical variables.

To explore a module network of interest and its associated meta-data in detail, users can click on a module name (Figure 1B). For example, Figure 2A displays a snapshot of the Angiogenesis enriched module network (module 26) within the ProNetView-ccRCC panel. Node sizes are proportional to the number of connections to let users easily identify the genes that are more central than others. To interactively explore the network, users can hover their mouse over a node, which will display the gene name of the phosphopeptide, and click on the node, which will display the associated gene-level meta-data in a separate panel (Figure 2B). As shown in Figure 2B, using ProNetView-ccRCC, users can quickly visualize the location of ERG in the network and display the connected nodes. As shown, ERG is directly connected to 11 genes within the same module and maps to 5 different peptides. In addition, users can click on the Heatmap link (Figure 2B), which will direct them to ProTrack tool,^[16]^ which provides a web-based interface to explore ccRCC multi-omics data through interactive heatmaps. To query whether other genes have similar association to tumor grade as ERG, users can easily construct a simple query for those genes (Figure 2C) by specifying the phenotype of interest (grade) and the FDR cutoff value (FDR= 20%). Genes that satisfy these metrics are listed in the text box and are highlighted in the network exploration panel in red (Figure 2D). Users can click the Reset button to return to the original network display.

**Figure 2.**
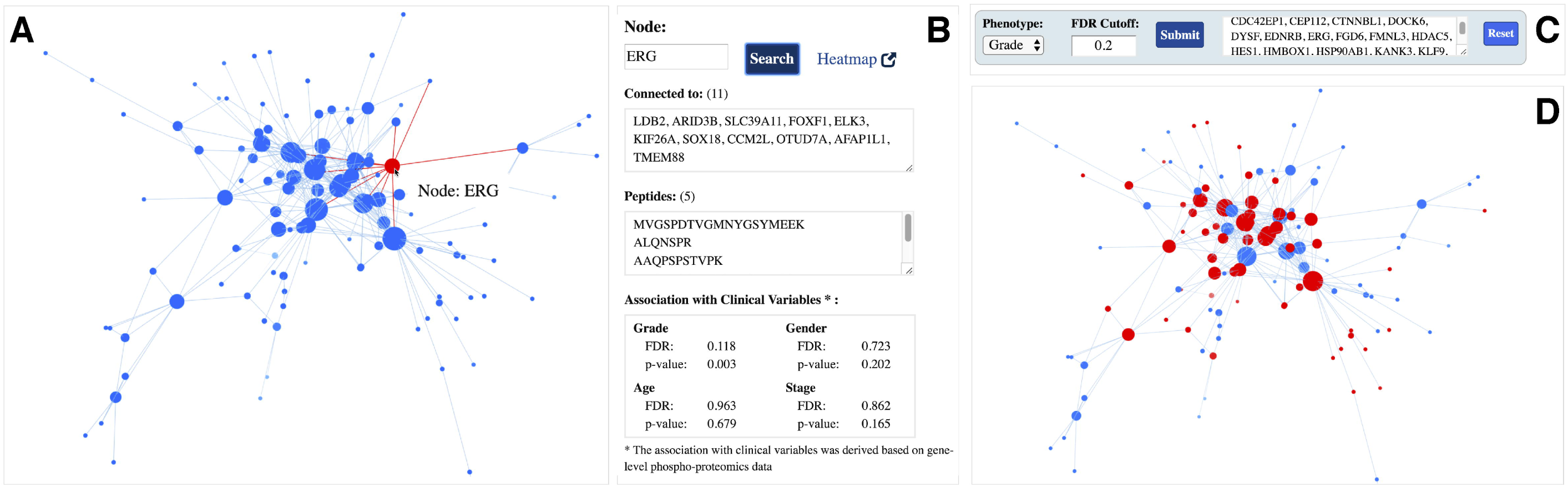
Angiogenesis enriched module network **A** interactive 3D network exploration panel. Hovering over gene ERG displays its name, highlights it in red, and clicking on the node highlights the connections in red as well; **B** meta-data information associated with ERG gene. Gene-specific information includes the list of connections (co-expressed genes), mapping peptides, and its association with clinical variables; **C** phenotype-based search panel. User selects the phenotype of interest and enters the FDR cutoff value. After clicking the Submit button, genes satisfying the criteria are listed in the text-box; **D** genes satisfying the phenotype-based search are highlighted in red in the interactive network exploration panel.

ProNetView-ccRCC also displays and provides interactive exploration capabilities for any pathway enriched in a network module. For example, Figure 3A displays the interactive table listing the enriched pathways, sorted by their p-values, associated with module 18. To identify the list of genes associated with a certain pathway (e.g. Cell cycle), users can click on the pathway name, which returns the list in a textbox (e.g. BRCA1, CDC20, CDK1, etc.) as well as highlights in the network exploration panel in red (Figure 3B). Clicking on the Reset button returns to the original network display.

**Figure 3.**
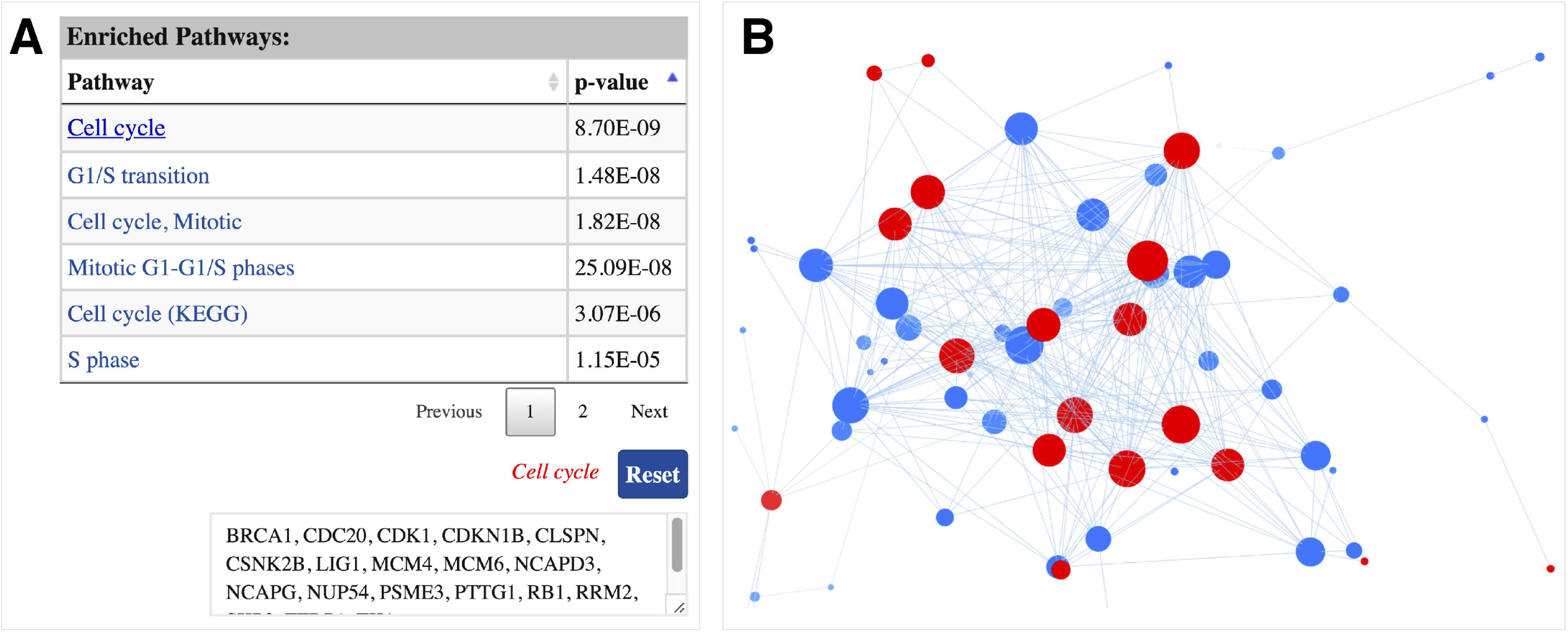
Cell cycle enriched module network **A** interactive table displaying the list of enriched pathways, sorted by their p-values, associated with the network. List of genes associated with a pathway can be displayed by clicking on the pathway name; **B** genes associated with the pathway of interest are highlighted in red in the interactive network exploration panel.

In conclusion, ProNetView-ccRCC, provides a free, open access and custom environment for the interactive exploration of CPTAC ccRCC networks and their associated meta-data. The tool runs on any modern web browser without the need for installing any specific plugins or libraries. We anticipate that ProNetView-ccRCC will facilitate researchers from a wide-spectrum computational skill levels to conduct their own analyses on the rich CPTAC ccRCC network data, and share and communicate their results.

## Abbreviations

CPTAC: Clinical Proteomic Tumor Analysis Consortium,
ccRCC: clear cell renal cell carcinoma;
iCAVE: interactome-CAVE

## ACKNOWLEDGEMENT

This work was supported in part by the NIH, National Cancer Institute’s Clinical Proteomic Tumor Analysis Consortium (CPTAC) grant U24CA210993.

